# “Validating silicon polytrodes with paired juxtacellular recordings: method and dataset”

**DOI:** 10.1101/037937

**Authors:** Joana P. Neto, Gonçalo Lopes, João Frazão, Joana Nogueira, Pedro Lacerda, Pedro Baião, Arno Aarts, Alexandru Andrei, Silke Musa, Elvira Fortunato, Pedro Barquinha, Adam R. Kampff

## Abstract

Cross-validating new methods for recording neural activity is necessary to accurately interpret and compare the signals they measure. Here we describe a procedure for precisely aligning two probes for *in vivo* “paired-recordings” such that the spiking activity of a single neuron is monitored with both a dense extracellular silicon polytrode and a juxtacellular micro-pipette. Our new method allows for efficient, reliable, and automated guidance of both probes to the same neural structure with micron resolution. We also describe a new dataset of paired-recordings, which is available online. We propose that our novel targeting system, and ever expanding cross-validation dataset, will be vital to the development of new algorithms for automatically detecting/sorting single-units, characterizing new electrode materials/designs, and resolving nagging questions regarding the origin and nature of extracellular neural signals.

## Introduction

Understanding how the brain works will require tools capable of measuring neural activity at a network scale, i.e., recording from thousands of individual neurons (Buzsáki, 2004). Technical advances have driven progress in large-scale neural recordings, and the development of microfabricated silicon polytrodes has led to an exponential increase in the number of neurons that can be simultaneously monitored (Stevenson and Kording, 2011)(Berényi et al., 2014)(Michon et al., 2014)(Nicolelis Lab, 2012). However, each improvement in recording technology inevitably raises new questions about the nature of the signal and demands new analysis methods to interpret these growing datasets.

Extracellular recording is unique in its ability to record populations of neurons deep in the brain with sub-millisecond resolution; it also poses particularly daunting challenges for analysis. Each electrode is sensitive to the spiking activity of hundreds of neurons in its vicinity, and sorting this cacophony into individual sources is a challenge (Marblestone et al., 2013). Furthermore, fundamental questions regarding how each neuron participates in the bulk extracellular signal remain unresolved: How many neurons contribute to the signal detected by an electrode? How does a neuron’s contribution decay with distance from the probe? Do different types of neurons have different extracellular signatures? Are extracellular recordings biased for particular types of neurons? How does the presence of the probe interfere with the activity of the surrounding neural tissue? Answers to these questions will require experiments to validate existing and future extracellular electrode technology as well as new analysis methods to interpret their data.

Employing modern methods for integrated circuit design and fabrication, probes with thousands, or even millions, of discrete sites are now being developed (Dombovári et al., 2014)(Ruther and Paul, 2015)(Shobe et al., 2015). These devices will densely sample the extracellular electric field, such that one nearby neuron will be detected by many individual electrodes, and will thus provide a detailed description of the spatiotemporal profile of a neuron’s extracellular action potential. It is expected that this additional detail will significantly aid analysis methods for the detection and isolation, and possibly type identification, of individual neurons in the vicinity of the probe, yet methods capable of utilizing such a dense sampling are just now being developed (Rossant et al., 2015).

“Ground-truth” data, for which one knows exactly when a neuron in the vicinity of an extracellular probe generates an action potential, is necessary to validate the performance of these new recording devices and analysis procedures. However, the validation datasets currently available for extracellular recordings only exist for tetrodes and single-wire electrodes (Wehr et al., 1999)(Henze et al., 2000) or are from *in vitro* preparations (e.g. slices (Anastassiou et al., 2015)) in which the majority of background neural activity has been surgically removed. Evaluating the existing silicon polytrodes, as well as forthcoming ultra-high density CMOS probes, *in vivo* will require new datasets, and, ideally, new methods for efficiently gathering this vital cross-validation data.

Targeting a single neuron close to an extracellular probe with another electrode requires accurately positioning both devices deep in neural tissue. When performed blindly, the efficiency of achieving “paired recordings” in which one neuron is detected by both probes is rather low, making such validation experiments much more difficult than just haphazardly recording extracellular neural signals. For this reason, such datasets are very rare, however, the ones that do exist (for tetrodes in the hippocampus) have been invaluable (Harris et al., 2000)(Gold et al., 2007). We anticipate that a large amount of such validation data will be required to characterize the large-scale neural recording devices currently being developed, and we thus set out to make paired recordings easier.

In the following we report a new method for efficiently and reliably targeting, blindly, two different recording devices to the same region in the brain. This method was then used to acquire a “ground truth” dataset from rat cortex with 32 and 128-channel silicon polytrodes, which can now be used to validate methods for interpreting dense extracellular recordings and help resolve persistent debates about the nature and origin of the extracellular signal. This dataset, which will grow as new devices are fabricated, is available online (http://www.kampff-lab.org/validating-electrodes).

## Materials and Methods

### Set-up design and calibration

The dual-recording setup requires two aligned, multi-axis micromanipulators (Scientifica, UK) and a long working distance optical microscope (Figure 1a). A “PatchStar” (PS) and an “In-Vivo Manipulator” (IVM) are mounted on opposite sides of a rodent stereotaxic frame. The probes were held at an angle: the PS allows the combination of two motion axes (XZ axis in approach mode) and this approach angle was set at 61° from the horizontal, whereas the IVM, a rigid 3-axis linear actuator, was tilted −48.2° from the horizontal around the Y axis. The use of two different models of manipulator was a practical constraint in this study, as the IVM permitted a greater range of travel for initial prototyping. However, future dual-probe setups will utilize two PS systems and the calibration and operation procedures will remain identical.

**Figure 1:**
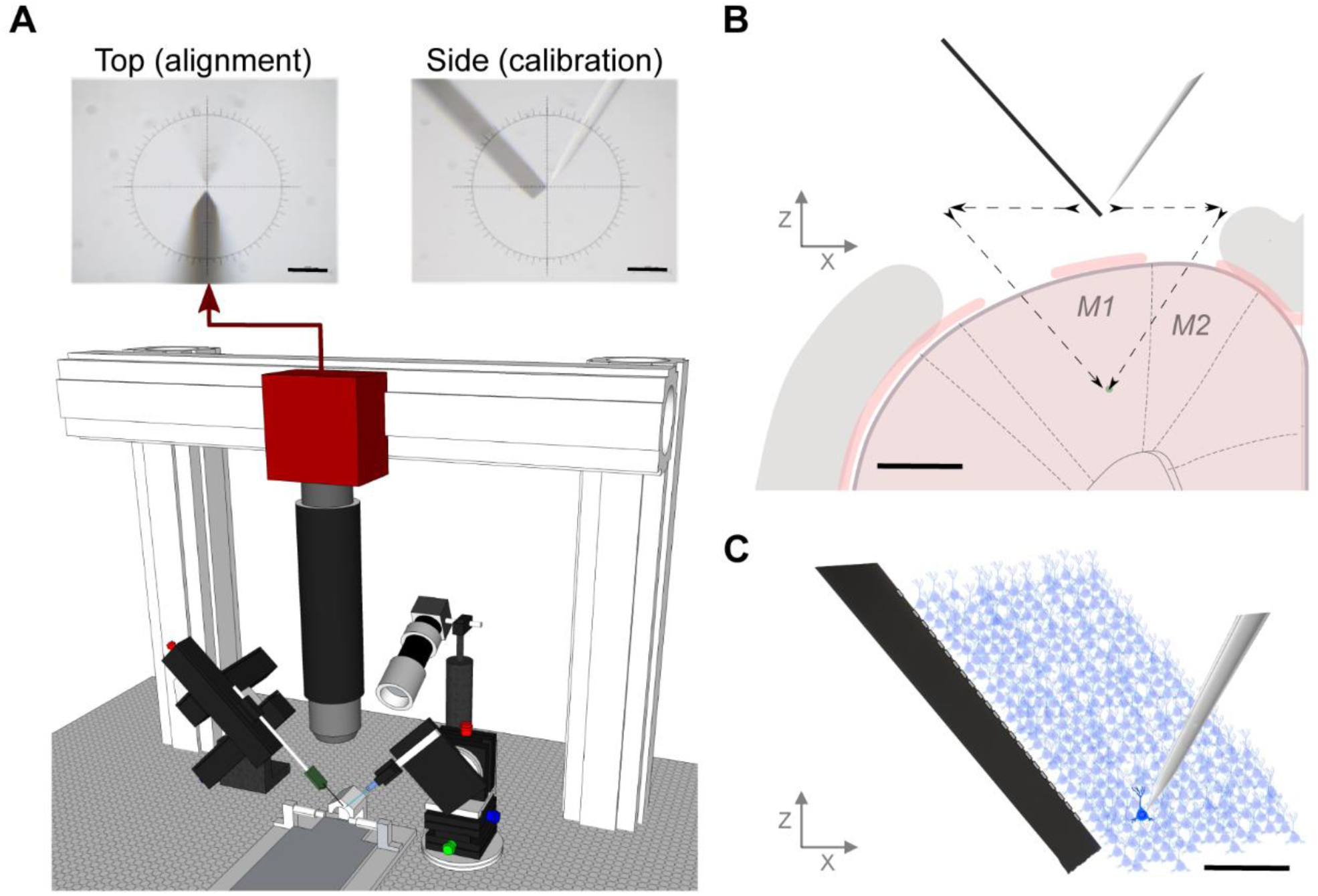
*In vivo* paired-recording setup: design and method. (a) Schematic of the dual-probe recording station. The PS micromanipulator drives the juxtacellular pipette and the IVM manipulator drives the extracellular polytrode. The setup includes a long working distance microscope assembled from optomechanical components mounted on a three-axis motorized stage. The alignment image provides a high-resolution view from above the stereotactic frame, upper left, however a side-view can also be obtained for calibration purposes, upper right (scale bar 100 μm). (b) Schematic of a coronal view of the craniotomy and durotomies with both probes positioned at the calibration point. The distance between durotomies, such that the probe tips meet at layer 5 in cortex, was around 2 mm. The black arrows represent the motion path for both electrodes entering the brain (scale bar 1mm). (c) Diagram of simultaneous extracellular and juxtacellular paired-recording of the same neuron at a distance of 90 μm between the micropipette tip and the closest electrode on the extracellular polytrode (scale bar 100 μm).

#### Alignment Microscope

A custom microscope was assembled from optomechanic components (Thorlabs, USA), a long-working distance objective (Infinity-Corrected Long Working Distance Objective, 10×, Mitutoyo, Japan), and a high-resolution CMOS camera (PointGrey, CA). The numerical aperture of the objective (0.28) had a theoretical resolution limit of ∼1 μm in X and Y directions and ∼10 μm in Z direction, which was oversampled by the camera sensor. Oblique illumination was necessary to acquire an image of both the extracellular probe and juxtacellular pipette, directly above the craniotomy, with sufficient contrast to accurately “zero” the position of each probe (Figure 1a, upper left panel). The repeatability of visually aligning each probe to the center of the image (“zeroing”) was evaluated by manually moving the tip of the pipette several times (n = 11) from outside the field-of-view to the focal plane and image center and recording the manipulator coordinates. The optical alignment procedure had 0.5 ± 0.5 μm repeatability in XY and 2.6 ± 1.7 μm in Z.

#### Mechanical calibration

Ensuring that the axes from both manipulators are parallel involved first a mechanical calibration procedure. The PS was aligned (“squared”) with the stereotactic frame using a machinist dial in exactly the same way that a machinist squares a milling machine XY table and column. The IVM manipulator was aligned using the same method in the Y axis using both the vertical and horizontal planes of the stereotaxic frame.

#### Software calibration

For each manipulator the position values are provided by the Scientifica software. To have both manipulators in the same reference frame it is necessary to apply a coordinate transformation to compensate for the angle difference between their respective motion axis. In the simplest case, this coordinate transformation would just use the angle measured by a gauge (Θ_IVM_ = 48.2°) and apply the following transformation:

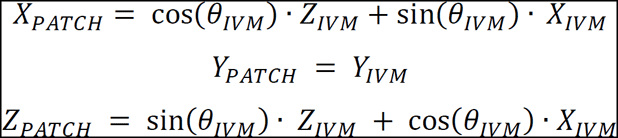

This transformation assumes that the Y axis of both manipulators are perfectly aligned. To obtain a more precise calibration, we adopted the PS coordinate system as the reference frame and transformed the recorded IVM coordinates into the PS frame, in an affine manner, as follows for the X axis: X_PATCH_ = A·X_IVM_, where A represents a transformation matrix that best matches these pairs of coordinates. In order to estimate this matrix, we moved the juxtacellular pipette to different points in space and then moved the extracellular electrode tip to the same points in space. The coordinates of each probe were recorded during this procedure using the optical microscope in a side-view “calibration” configuration (Figure 1a, upper right panel). These measurements were repeated several times at several locations (n = 20) so that an appreciable volume (400 × 600 × 1100 μm) was canvassed. The protocol for the acquisition and transformation of axis position was implemented in Bonsai, an open-source visual programming framework, which can be freely downloaded at https://bitbucket.org/horizongir/bonsai (Lopes et al., 2015).

### Surgery

Rats (400 to 700 g, both sexes) of the Long-Evans strain were anesthetized with a mixture of Ketamine (60 mg/kg intraperitoneal, IP) and Medetomidine (0.5 mg/kg IP) and placed in a stereotaxic frame that was atop a vibration isolation table (Newport, USA). Equipment for monitoring body temperature as well as a live video system for performing craniotomies and durotomies was integrated into the setup. At the initial stage of each surgery atropine was given to suppress mucus secretion (Atropine methyl nitrate, Sigma-Aldrich). Anesthetized rodents then underwent a surgical procedure to remove the skin and expose the skull above the targeted brain region. Small craniotomies (4 mm medial-lateral and 2 mm anterior-posterior) were performed above dorsal cortex. The craniotomy centers were 2.5 mm lateral to the midline and ranged from +4 to −4 anterior-posterior, thus exposing either motor, sensory or parietal cortex. Two reference electrodes Ag-AgCl wires (Science Products GmbH, E-255) were inserted at the posterior part of the skin incision on opposite sides of the skull.

### Dense silicon polytrodes

All experiments were performed with two different high-density silicon polytrodes. A commercially available 32-channel probe (A1x32-Poly3-5mm-25s-177-CM32, NeuroNexus, USA), with 177 μm^2^ area electrodes (iridium) and an inter-site pitch of 22-25 μm, was used in the first experiments. The impedance magnitude for these sites at 1 kHz was ∼1 MΩ, but for some experiments, a PEDOT coating was applied to lower this impedance to ∼100 kΩ. In later experiments, we used a 128-channel probe conceived in the collaborative NeuroSeeker project (http://www.neuroseeker.eu/) and developed by IMEC using CMOS-compatible process technology. This probe’s TiN electrode were 400 μm^2^ (20 × 20 μm^2^) large, arranged at a pitch of 22.5 μm, and had an impedance magnitude of 50 kΩ at 1 kHz.

Before each surgery, the impedance magnitude of each electrode site was measured for diagnostic purposes using a protocol implemented by the amplifier/DAC chip (InTan Technologies, USA). Following each surgery, cleaning was performed by immersing the probe in a trypsin solution (Trypsin-EDTA (0.25%), phenol red, TermoFisher Scientific) for 30-120 minutes and rinsing with distilled water.

### Probe insertion and simultaneous juxtacellular-extracellular recordings

After both the extracellular probe and juxtacellular pipette positions were sequentially “zeroed” to the center of the microscope image, the microscope was replaced by a macro-zoom lens (Edmund Optics, USA) for visually guided insertion. The extracellular probe was inserted first, at a constant velocity of 1 μms^-1^, automatically controlled by the manipulator software. When the extracellular probe was in place, the juxtacellular pipette, pulled from 1.5 mm capillary borosilicate glass (Warner Instruments, USA) and filled with PBS 1×, was then lowered through a second durotomy under visual guidance using the overhead surgery camera.

As the pipette approached the extracellular electrodes, we followed a protocol for performing loose-patch recordings from neurons as previously described (Herfst et al., 2012). Positive pressure (25-30 mmHg) was reduced on the pipette to 1-10 mmHg (DPM1B Pneumatic Transducer Tester, Fluke Biomedical, USA) and the amplifier for juxtacellular recordings (ELC-01X, NPI, Germany) was set to voltage-clamp mode (25 mV steps at 20Hz). As the electrode was advanced towards a cell membrane, we observed an increase in the pipette resistance. If spikes were observed, the pressure was then released (0 mmHg) and a slight suction applied to obtain a stable attachment to the cell membrane.

A data acquisition board (National Instruments, USA) was used to control amplifier voltage commands. However, after a stable recording is achieved, simultaneous recording of both extracellular and juxtacellular electrodes used exclusively the Open Ephys (http://www.open-ephys.org) acquisition board ADCs (for the juxtacellular signal) along with the RHD2000 series digital electrophysiology interface chip that amplifies and digitally multiplexes the extracellular electrodes (Intan Technologies, USA). Extracellular signals in a frequency band of 0.1-7500 Hz and juxtacellular signals in a frequency band of 300-8000 Hz were sampled at 30 kHz with 16-bit resolution and were saved in a raw binary format for subsequent offline analysis using a Bonsai interface. For the analyses described in the following, a 3rd order Butterworth filter with a band-pass of 100-14250 Hz (95% of the Nyquist frequency) was used in the forward-backward mode. For some recordings we noticed a high-frequency noise contribution and we thus used a band-pass of 100-5000 Hz.

All experiments were approved by the Champalimaud Foundation Bioethics Committee and the Portuguese National Authority for Animal Health, Direcção-Geral de Alimentação e Veterinària (DGAV).

## Results

### Setup design

The “dual-probe” positioning and recording setup presented in Figure 1a was designed to reliably target neural cell bodies located within ∼100 μm of the polytrode electrode sites without optical guidance. In this setup, the motorized manipulators, video capture, online visualization/control parameters, and extra- and juxtacellular voltage recording were integrated and coordinated by custom open-source software developed within the Bonsai framework (Lopes et al., 2015).

Following a mechanical alignment and software calibration of both manipulators’ axes, each paired-recording experiment began with the optical “zeroing” of both probes. Each probe was positioned, sequentially, at the center of the microscope image (indicated by a crosshair) and the motorized manipulator coordinates set to zero (Figure 1a). As shown in Figure 1b, this alignment is performed directly above the desired *rendez-vous* point inside the brain, as close as possible above dura, usually between 1 and 4 mm, but far enough to reduce background light reflected from the brain surface into the microscope image. During optical calibration it is possible to select any point on the extracellular electrode to be the origin (X=0, Y=0, Z=0) by aligning that point of the probe in the reticle. However, the distance reported in the subsequent data is always the Euclidean distance between the tip of the pipette and the closest extracellular electrode.

With practice, multiple cells in the vicinity (less than 200 μm) of the polytrode could be recorded during a single insertion of the juxtacellular pipette (Figure 1c is a schematized example of one such paired recording, *2014_03_26_Pair2.0*).

Prior to the surgeries, we validated the alignment of the motors by moving both probes independently from the calibration point (0, 0, 0) towards a different point in space and recording the position difference between them after travel (see Methods). During our experiments, the movement of the probes primarily occurred in the XZ plane. We found that when we moved both probes to a new Z position 3 mm below the calibration point (0, 0, −3), similar to an actual recording experiment (Figure 1b), the position error observed was reliably less than 30 μm after the software calibration (and 40 μm before software calibration), which was acceptable for targeting the same region in cortex.

### Paired juxtacellular-extracellular recording

Twenty-three paired recordings with a distance less than 200 μm between the juxtacellular neuron and closest extracellular electrode were obtained from the cortex of anesthetized rats. The juxtacellular pipette had a long thin taper that enables minimal tissue damage and displacement during penetration (Figure 2a). As the juxtacellular electrode was advanced through the brain, several neurons were encountered at different locations along the motion path and, consequently, at different distances from the extracellular polytrodes. Figure 2b illustrates the large juxtacellular (peak-to-peak ∼4 mV) signal recorded from a neuron encountered at a distance of 68 ± 30 μm between the micropipette tip and the closest extracellular electrode. The positive-before-negative biphasic waveform shape (Figure 2c) is indicative of a capacitively-coupled cell-attached recording from a somatic/perisomatic located recording pipette (Herfst et al., 2012). However, two juxtacellular recordings in the dataset exhibit the waveform of well isolated extracellular spike (negative-before-positive), likely due to incomplete contact between the membrane and pipette presenting lower peak-to-peak amplitudes (*2015_09_04_Pair 5.0* and *2015_09_03_Pair 9.0*).

**Figure 2:**
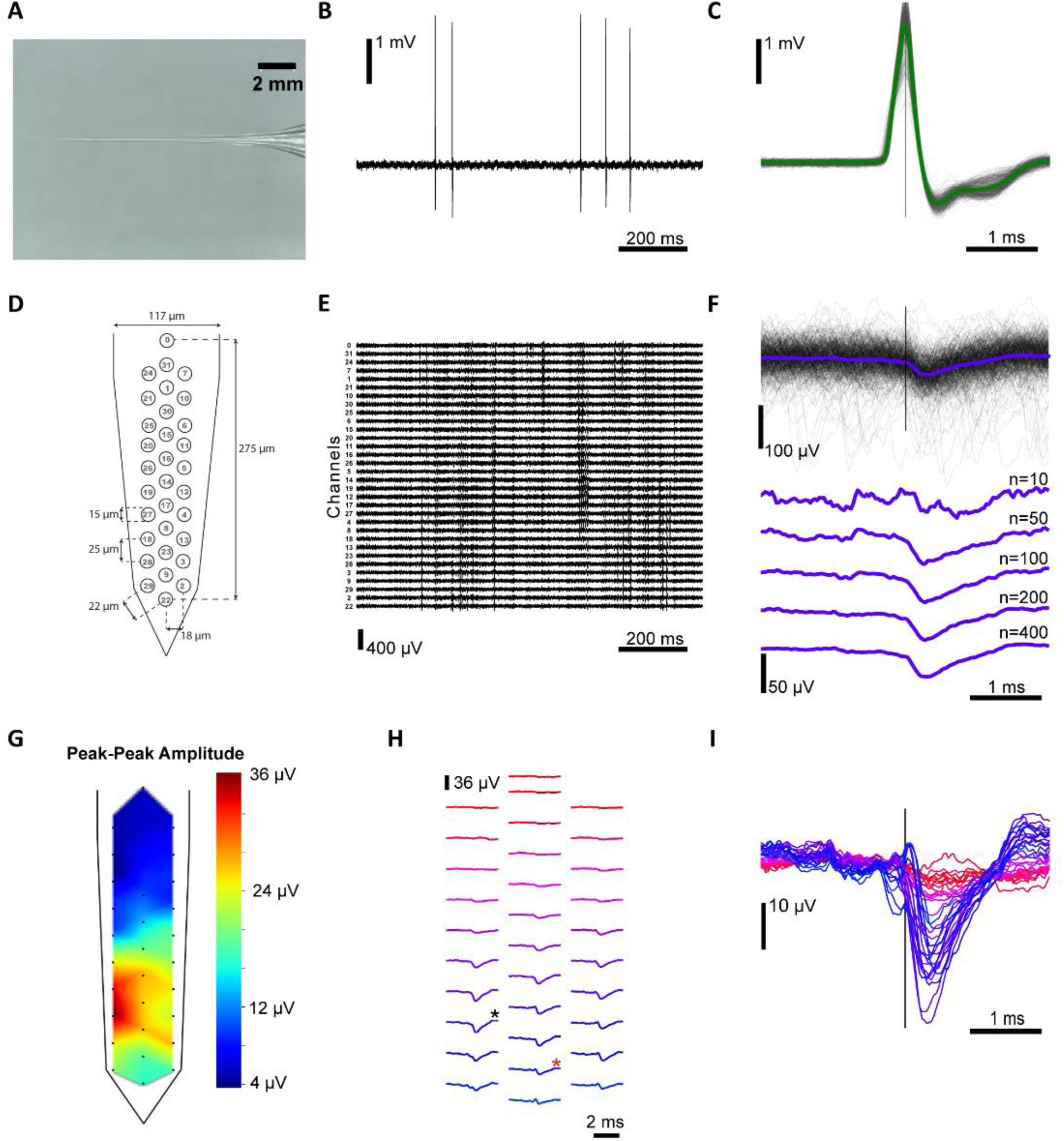
Paired extracellular and juxtacellular recordings from the same neuron. (a) Recording pipette with a long thin taper used for juxtacellular recordings with typical tip diameter of 1-4 μm and resistance of 3-7 MQ. (b) Representative juxtacellular recording from a cell in layer 5 of motor cortex, 68 μm from the extracellular probe *(2014_10_17_Pair1.0)*, with a firing rate of 0.9 Hz. (c) The juxtacellular action potentials are overlaid, time-locked to the maximum positive peak, with the average spike waveform superimposed (n= 442 spikes). (d) Extracellular dense polytrode array with an inter-site pitch of 22-25 μm and a span of 275 μm along the shank axis; the electrode channel number is represented at each site. (e) Representative extracellular recording that corresponds to the same time window as the above juxtacellular recording. Traces are ordered from upper to lower electrodes and channel numbers are indicated. (f) Extracellular waveforms, aligned on the juxtacellular spike peak, for a single channel (channel 18) and the juxtacellular triggered average (JTA) obtained by including an increasing number of juxtacellular events (n as indicated). (g) Spatial distribution of the amplitude for each channel’s extracellular JTA waveform. The peak-to-peak amplitude within a time window (+/- 1 ms) surrounding the juxtacellular event was measured and the indicated color code was used to display and interpolate these amplitudes throughout the probe shaft. (h) The waveform averages for all the extracellular electrodes are spatially arranged. The channel with the highest peak-to-peak JTA (channel 18) is marked with a black (*) and the closest channel (channel 9) is marked with a red (*). (i) The extracellular JTA time courses for each channel are overlaid and colored according to the scheme in (h).

A simultaneous extracellular recording was made with the 32-channel probe illustrated in Figure 2d, allowing us to specifically characterize the extracellular signature of an action potential generated by the juxtacellular recorded neuron. The band-pass filtered extracellular traces, ordered according to the electrode’s geometry, are presented in Figure 2e and correspond to the same time window as the juxtacellular recording (Figure 2b). A short time window (4 ms) extracted from the extracellular trace around each detected juxtacellular event (occurrence of the action potential positive peak) for one extracellular channel is shown in Figure 2f. Despite the low amplitude, a clear extracellular signature of the juxtacellular recorded neuron’s spike can be recovered by averaging windows across multiple events. This juxtacellular triggered average (JTA) can be computed for all channels, allowing a high signal-to-noise estimate of the spatiotemporal distribution of the extracellular action potential. The JTA peak-to-peak amplitude for each channel interpolated within the electrode site geometry, sometimes called “the cell footprint” (Delgado Ruz and Schultz, 2014), is shown in Figure 2g. The JTA waveforms for each channel are shown, arranged using the relative probe spacing in Figure 2h and overlaid in Figure 2i.

The example presented in Figure 2 is from one paired juxtacellular and extracellular recording. Several recordings were made in a similar manner and we next examined the variety of extracellular signatures obtained for different neurons at different positions relative to 32 and 128-channel dense polytrodes.

#### Distance dependence of extracellular signal amplitude

Across all of our paired-recordings (Figure 3) the distance between a neuron and the extracellular electrodes was the major factor determining the peak extracellular signal amplitude. For neurons encountered within 11 to 180 μm from the probe surface, the magnitude of the neuron’s extracellular signal ranges from 416 to 7 μV. In our dataset, the closest neurons (< 60 μm) had very strong signals on the extracellular probe. However, neurons ∼100 μm from the probe exhibited a broad range of extracellular signal peak amplitudes (from ∼7-60 μV). At larger distances we couldn’t distinguish a clear spike waveform from the small (>1 μV) cross-talk artifact even after averaging. Nevertheless, we include these cells in the dataset since they could potentially be used to better understand spike-LFP relationships (Lewis et al., 2015)(Berényi et al., 2014).

**Figure 3:**
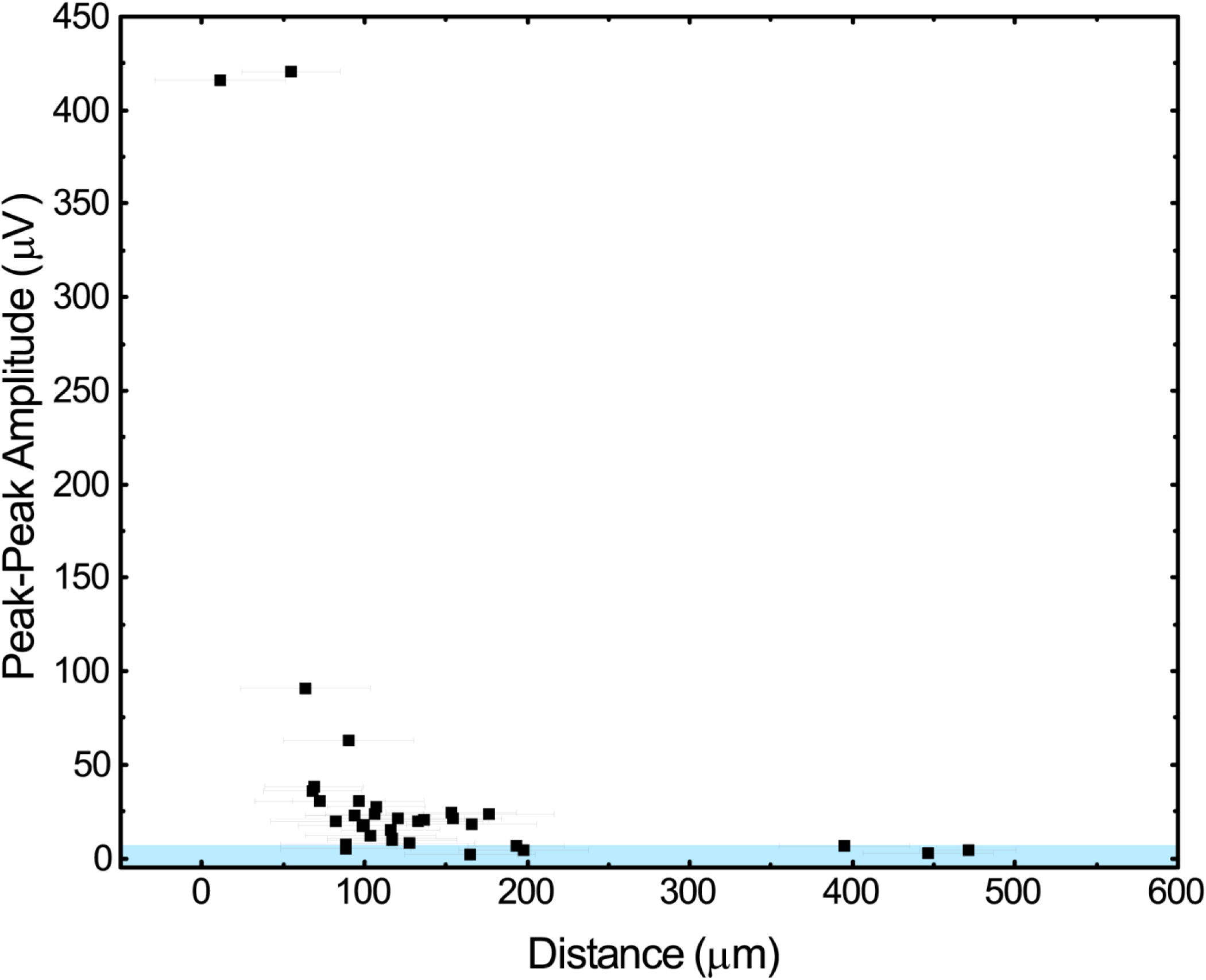
Distance dependence of extracellular signal amplitude. The maximum peak-to-peak amplitude of the JTAs (+/- 1 ms of the alignment time) across all extracellular channels for each paired recording vs. the distance between the closest extracellular electrode and the juxtacellular pipette tip. Horizontal error bars report uncertainty in position estimate (+/- 30 or 40 μm). The blue shaded region indicates a 5 μV threshold for excluding possible cross-talk electrical artifacts between the extra- and juxtacellular recording electronics.

#### Detection of the juxtacellular spikes on the extracellular probe

The first step in the analysis of extracellular data is the identification of discrete spike events (Hazan et al., 2006). Therefore, in order to use paired recordings to evaluate algorithms for assigning these extracellular events to clusters belonging to distinct neurons (i.e spike sorting), one must be able to detect the juxtacellular spike on the extracellular electrodes. We used a popular spike detection algorithm, SpikeDetekt, which extracts action potentials as spatiotemporally localized events (Rossant et al., 2015), to identify all spikes visible to our extracellular probe and then looked for correspondences with the juxtacellular spike times. The SpikeDetekt algorithm was used with high-pass filtered data (3rd order Butterworth, forward-backward mode, band-pass filter of 500-14250 Hz) and a strong and weak threshold of 4.5 and 2 times the standard deviation, respectively. The juxtacellular spike times were determined as the peaks of well-isolated threshold crossings (Supplementary Table 1).

This spike detection process is illustrated in Figure 4a-c for a data segment containing the contribution of a neuron that was simultaneously recorded with the juxtacellular pipette (*2014_11_25_Pair3.0*). To compare the extracellularly detected event times with the spike times observed in the juxtacellular recording, we generated a peri-event time histogram (PETH) using all spike events found on the extracellular channels aligned relative to each juxtacellular spike (Figure 4 d, e). In some paired recording datasets, these PETHs reveal a high probability of spike co-occurrence at 0 ms, indicating that the juxtacellular neuron’s spike is being found by SpikeDetekt. The count value in the PETH 0 ms bin for the recording pair in Figure 4d *(2014_11_25_pair3.0, channel 18)* suggests that all of the juxtacellular spikes were found, but that this bin also included detections of coincident spiking events occurring in the background neural activity (386 detected, 348 actual). In contrast, the PETH 0 ms bin for the pair in Figure 4e *(2014_ 03_26_pair2.0, channel 9)* indicates that only a fraction of the total number of juxtacellular spikes were detected (35 detected, 150 actual). The low amplitude of the extracellular spike (the magnitude of that neuron’s signal is closer to the electrode’s noise level) in Figure 4e is a challenge for the single channel threshold used by SpikeDetekt.

**Figure 4:**
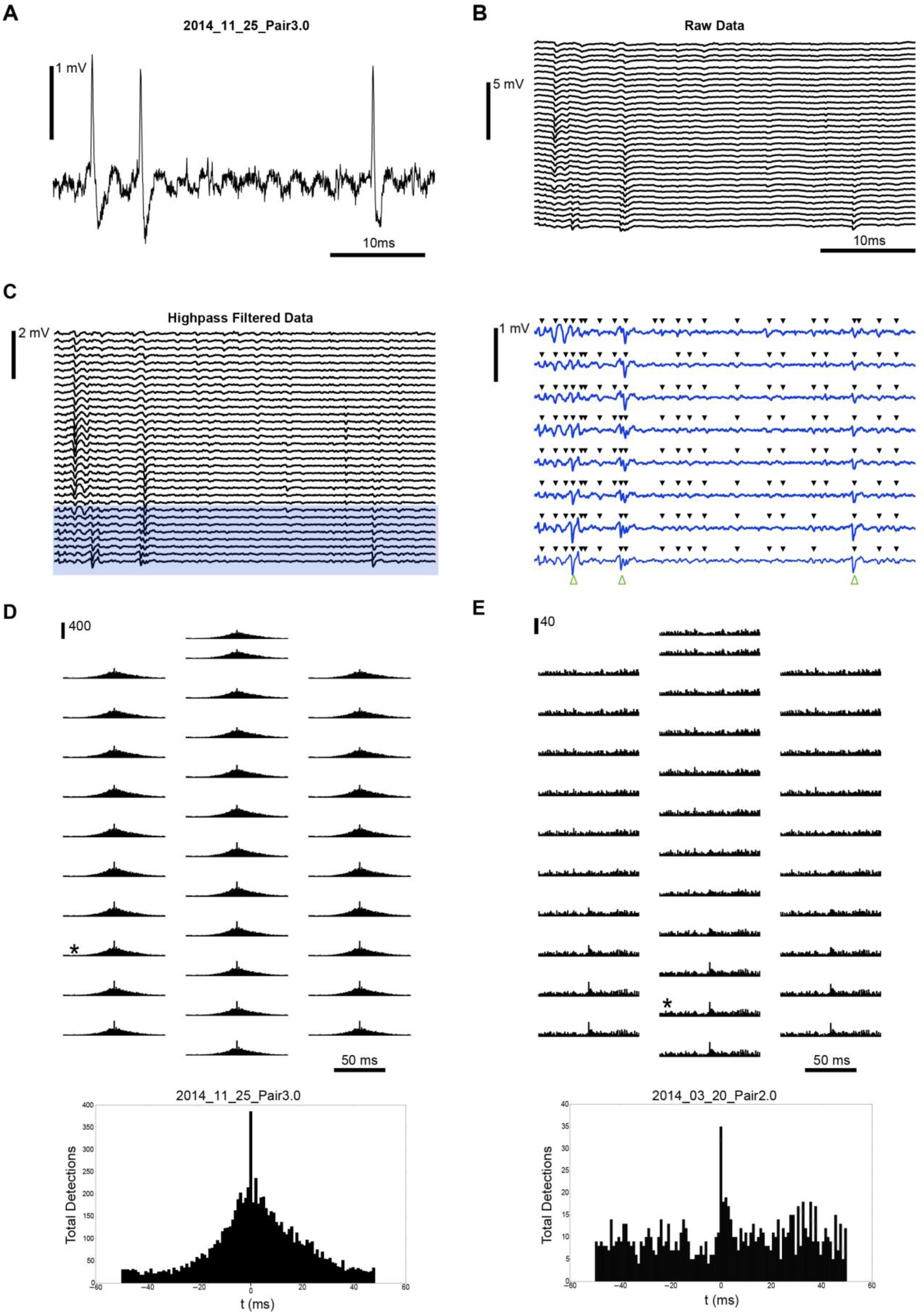
Extracellular detection of the juxtacellular neuron’s actional potentials. (a) Representative juxtacellular recording and (b) wide-band (0.1-7500 kHz) signal recorded simultaneously with a 32-channel silicon polytrode. (c) The extracellular data after highpass filtering (500 Hz), green arrows indicate the time of juxtacellular spikes (left panel). The highlighted traces are expanded in the right panel and included black arrows to indicate all spikes identified by SpikeDetekt using standard thresholds. (d) Peri-event time histograms of the extracellular spike events found by SpikeDetekt, relative to the juxtacellular spike times in 1 ms bins centered at 0 ms, are shown for each electrode channel at their relative position on the extracellular probe. The channel with the largest peak in the bin at 0 (+/- 0.5 ms from the juxtacellular event) is indicated (*) and expanded in the lower panel. (e) The same presentation as in (d), but for a neuron with a smaller extracellular action potential.

#### Spatiotemporal structure of extracellular signatures

Neurons near the polytrode surface exhibited a rich diversity of action potential waveforms (amplitude and dynamics) spread across multiple electrode sites (Figure 2h, Figure 5 b, e and Figure 6). This spatiotemporal structure will not only provide additional information for improving spike detection and sorting procedures, but may also reveal specific contributions from different parts of the neural membrane to this extracellular signature (Hu et al., 2009). For example, in Figure 5c, after hypothesizing a relative position and morphology for the recorded neuron (Figure 5a), the first negative peak (bottom blue trace) in the extracellular potential may arise from currents in the distal axon initial segment and the later peaks might then be due to the backpropagation to the soma (middle, purple trace) and on to the dendrites (top red trace). Another example of complex structure in the extracellular signature is seen in Figure 5e, in this case the primary signal is localized to a small region of electrodes and varies greatly between neighboring sites, which are separated by only ∼20 μm. These, and other examples in our dataset, clearly suggest that the amount of useful spatiotemporal information captured by dense large-scale neural recording devices is promising (Buzsáki, 2004), not only for improving algorithms that detect and sort events, but also to identify cell-types based on the morphology suggested by their extracellular signature.

**Figure 5:**
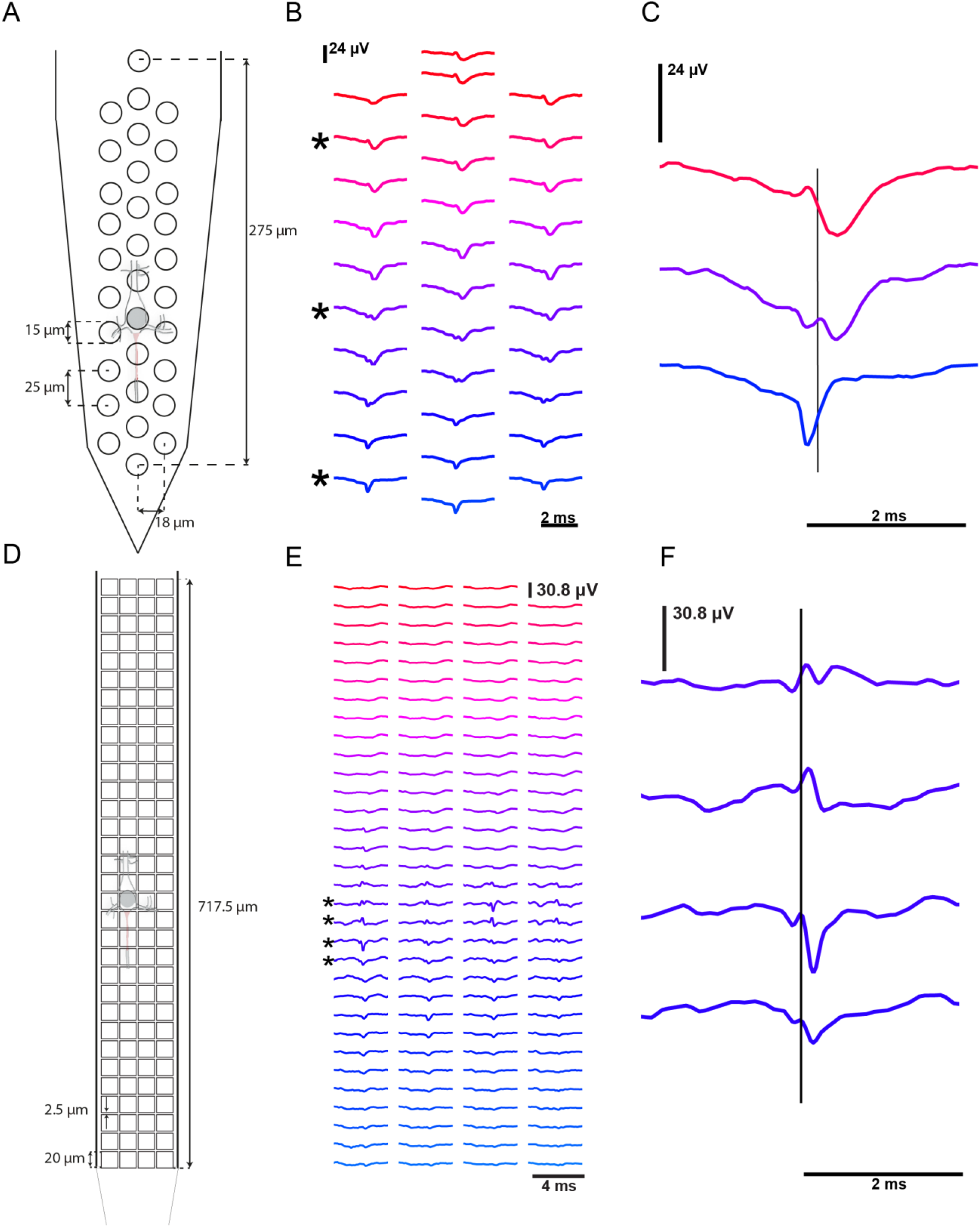
Spatiotemporal structure of extracellular signatures. (a) The hypothesized relationship between the 32-channel electrode array and a juxtacellular recorded neuron *(2014_11_25_Pair1.1)* is schematized; the axon initial segment is depicted in red. (b) The JTA waveforms for all the extracellular electrodes are spatially arranged according to the probe geometry. (c) Expanded comparison of the JTA waveforms for the indicated electrodes with a line denoting the peak time of the juxtacellular spike. (d-f) Similar presentation as (a-c) for the 128-channel polytrode *(2015_0S_04_Pair5.0)*.

**Figure 6:**
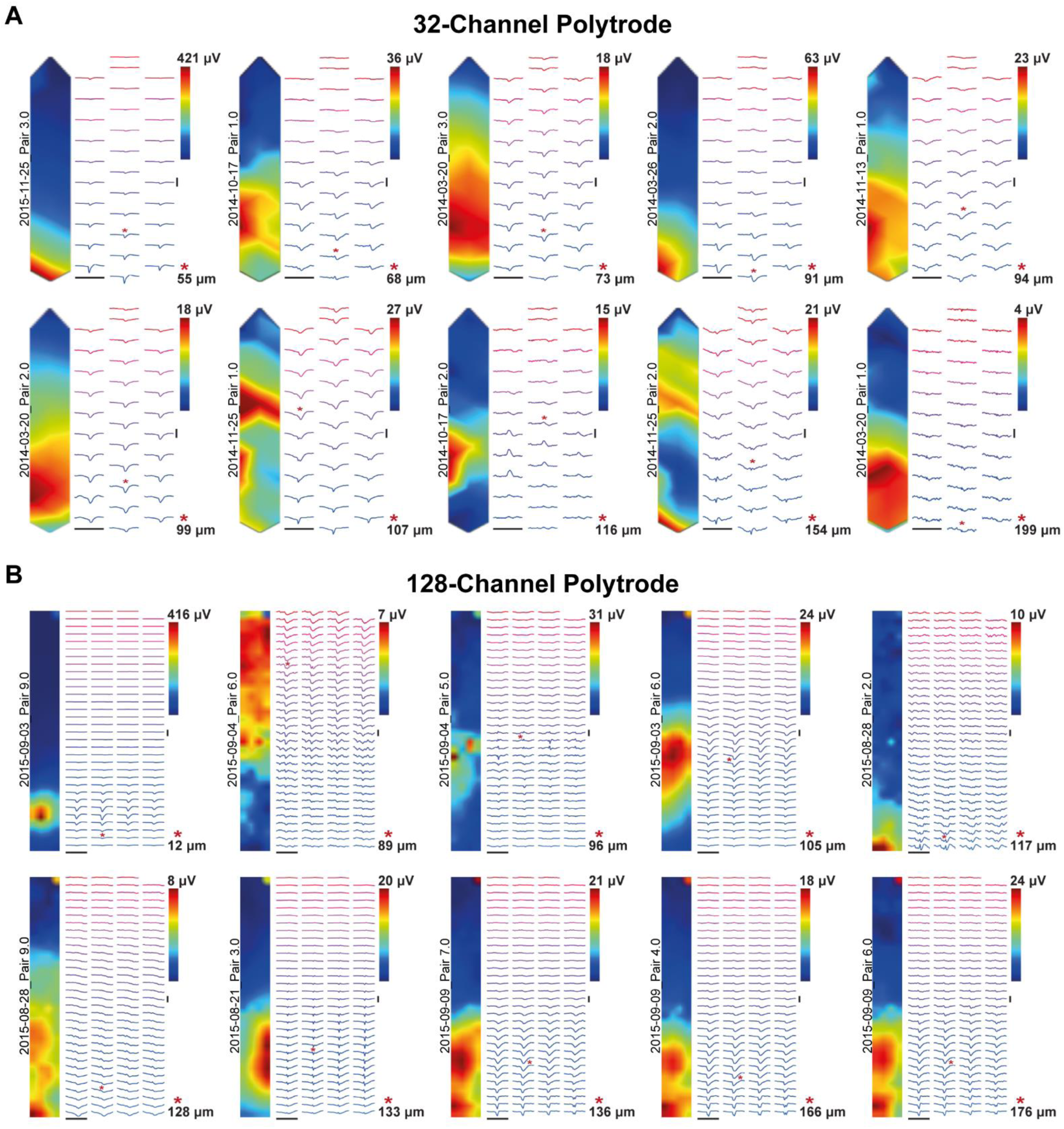
Dataset for validating spike detection and sorting algorithms for dense polytrodes. (a) Spatial distribution of the peak-to-peak amplitude within a time window (+/- 1 ms) surrounding the juxtacellular event and the indicated color code was used to display and interpolate these amplitudes throughout the 32-channel probe shaft. In addition, the extracellular JTA waveforms for all the extracellular electrodes are spatially arranged. (b) The same presentation as in (a) for paired-recordings with the 128-channel probe.

## Discussion

In the present study, our dual-recording setup allowed precise targeting of both an extracellular probe and a juxtacellular pipette to the same position in cortex. The setup is low-cost and easily implemented by any electrophysiology laboratory with two motorized (servo/stepper) micromanipulators. We hope that our description will guide/instigate the collection of such critical cross-validation data from the forthcoming deluge of novel neural recording devices.

### Dataset for cross-validating polytrodes and spike detection/sorting algorithms

A summary of the current cross-validation dataset is presented in Figure 6. It includes twenty juxta-extracellular pairs recorded with both 32 and 128-channel polytrodes, at a range of inter-probe distances, 800 to 1800 μm deep in cortex. The distance between the closest electrode and the neuron from which it recorded was critical in determining the amplitude of the extracellular spike (the maximum distance at which we observed a large peak-to-peak amplitude spike (> 50 μV) was 91 μm). However, for neurons recorded at a similar distance to each other (e.g. ∼100 μm) the signal amplitude, and the spatiotemporal signature, was quite variable (Figure 3 and 6). This variability may reflect the morphology of different neurons, heterogeneous properties of extracellular space (Bédard et al., 2004), the orientation of the neuron relative to the probe (Gold et al., 2006), and/or the extracellular signature of different cell types (Gray et al., 1995). Future experiments, which use cell-attached labeling to anatomically reconstruct the juxtacellular neuron following a paired recording, are now being pursued to resolve these open questions and extend the validation dataset. However, the existing dataset already includes some useful cross-validation examples: nearby cells with large extracellular action potentials, which will provide “ground-truth” data for evaluating current spike detection and sorting algorithms, as well as more “challenging” intermediate cells, for which new algorithms, specifically optimized to use the additional information available to dense silicon polytrodes, should be able to recover. The full dataset, as well as probe maps and analysis code, is available online (http://www.kampff-lab.org/validating-electrodes/) and summarized in Supplementary Table 1.

## Acknowledgements

This work was supported by funding from the European Union’s Seventh Framework Programme (FP7/2007-2013) under grant agreement nr. 600925 and the FCT-MCTES doctoral grants SFRH/BD/76004/2011 given to JN, and the Bial Foundation (Grant 190/12). Institutional support and funding was provided by the Champalimaud Foundation and Sainsbury Wellcome Centre (funded by the Gatsby Charitable Foundation and the Wellcome Trust). We are very grateful to George Dimitriadis, Kinga Kocsis and all the members of the Intelligent Systems Laboratory for their contributions. We also thank Kenneth Harris’ lab and members of the Bernstein Center in Freiburg for helpful discussions.

